# Tomato brown rugose fruit virus Mo gene is a novel microbial source tracking marker

**DOI:** 10.1101/2023.01.09.523366

**Authors:** Aravind Natarajan, Brayon J. Fremin, Danica T. Schmidtke, Marlene K. Wolfe, Soumaya Zlitni, Katherine E. Graham, Erin F. Brooks, Christopher J. Severyn, Kathleen M. Sakamoto, Norman J. Lacayo, Scott Kuersten, Jeff Koble, Glorianna Caves, Inna Kaplan, Upinder Singh, Prasanna Jagannathan, Andrew R. Rezvani, Ami S. Bhatt, Alexandria B. Boehm

## Abstract

Microbial source tracking (MST) identifies sources of fecal contamination in the environment using fecal host-associated markers. While there are numerous bacterial MST markers, there are few viral markers. Here we design and test novel viral MST markers based on tomato brown rugose fruit virus (ToBRFV) genomes. We assembled eight nearly complete genomes of ToBRFV from wastewater and stool samples from the San Francisco Bay Area in the United States of America. Next, we developed two novel probe-based RT-PCR assays based on conserved regions of the ToBRFV genome, and tested the markers’ sensitivities and specificities using human and non-human animal stool as well as wastewater. TheToBRFV markers are sensitive and specific; in human stool and wastewater, they are more prevalent and abundant than a currently used marker, the pepper mild mottle virus (PMMoV) coat protein (CP) gene. We applied the assays to detect fecal contamination in urban stormwater samples and found that the ToBRFV markers matched cross-assembly phage (crAssphage), an established viral MST marker, in prevalence across samples. Taken together, ToBRFV is a promising viral human-associated MST marker.

**Importance:** Human exposure to fecal contamination in the environment can cause transmission of infectious diseases. Microbial source tracking (MST) can identify sources of fecal contamination so that contamination can be remediated and human exposures can be reduced. MST requires the use of fecal host-associated MST markers. Here we design and test novel MST markers from genomes of tomato brown rugose fruit virus (ToBRFV). The markers are sensitive and specific to human stool, and highly abundant in human stool and wastewater samples.

## Introduction

Across the world, water quality is assessed for human fecal contamination using microbial indicators, including total coliforms like *Escherichia coli* and enterococci (1–3). Using these organisms to assess water quality is advantageous because they are abundant in human stool, which enables detection of even trace contamination of waterbodies. Additionally, their presence may indicate the potential contamination of waterbodies by other sparser human pathogens that may be harder to detect. However, there are limitations to their utility. These microbial indicators of human fecal contamination are also found in non-human stool (4). Additionally, they can be present and even grow in the environment, including in decaying plant material (1, 5), and in soils and sands (6, 7). Therefore, there is a need to identify new microbial indicator targets that can be used to specifically assess the presence of human fecal contamination.

The process of detecting microbes and identifying sources of microbial contamination in the environment is known as Microbial source tracking (MST). MST targets have also been used in COVID-19 wastewater-based epidemiology applications as “fecal strength” and endogenous extraction controls (8). Over the last decade, sensitive and specific molecular MST markers have been developed for various animal stools including human (9), cow (10), and birds (11). Most of these MST markers target conserved regions of bacterial genomes (9), with the exception of two that target the cross-assembly phage (crAssphage) (12) and pepper mild mottle virus (PMMoV) (13). crAssphage, a phage of *Bacteroidetes*, is a DNA virus that is highly abundant in the human gut (14). PMMoV is a plant RNA virus found at high concentrations in the human gut given its presence in popular spices, hot sauces, and other food products (15). The performance of MST targets is evaluated in terms of sensitivity and specificity for the host stool. For instance, a sensitive target for human stool is present at high concentrations in nearly all human fecal samples so that dilute human stool can be detected in the environment. Meanwhile, a specific target is absent in nearly all non-human fecal samples. A previous study defined an MST assay as being sensitive and specific if the true positive and true negative rates were greater than 80% (9).

In this study, we present a new human-associated, RNA-based, viral MST target that is highly abundant in human stool and wastewater, tomato brown rugose fruit virus (ToBRFV). ToBRFV was first identified in Israel in 2014 and has since been detected across the world. To date, ToBRFV has been found across four continents, in at least 35 countries; this is likely an underestimate (16). We assembled eight nearly complete genomes of ToBRFV from wastewater and stool samples from the San Francisco Bay Area (Bay Area) in California in the United States of America (U.S.A), representing some of the first complete genomes from stool and wastewater in the area. Using these complete genomes, and other publicly available genomes, we developed two novel hydrolysis probe-based RT-PCR assays based on conserved regions of its RNA genome, and tested their sensitivity and specificity using stool and wastewater samples. Finally, we apply this assay for MST in stormwater samples collected from an urban environment. With the finding that ToBRFV is a reliable RNA-virus based MST marker, this study makes a valuable contribution to detecting human fecal contamination of the environment, and to waste-water based epidemiology.

## Materials and methods

### Assembly and analysis of ToBRFV genomes, and design of hydrolysis-probe RT-PCR assays

In order to design ToBRFV-specific primers and probes for hydrolysis-probe RT-PCR assays, all ToBRFV genomes available in February 2021 were obtained. These were supplemented with new genomes assembled from stool samples processed and sequenced in this study (Table 1).

**Table 1:**
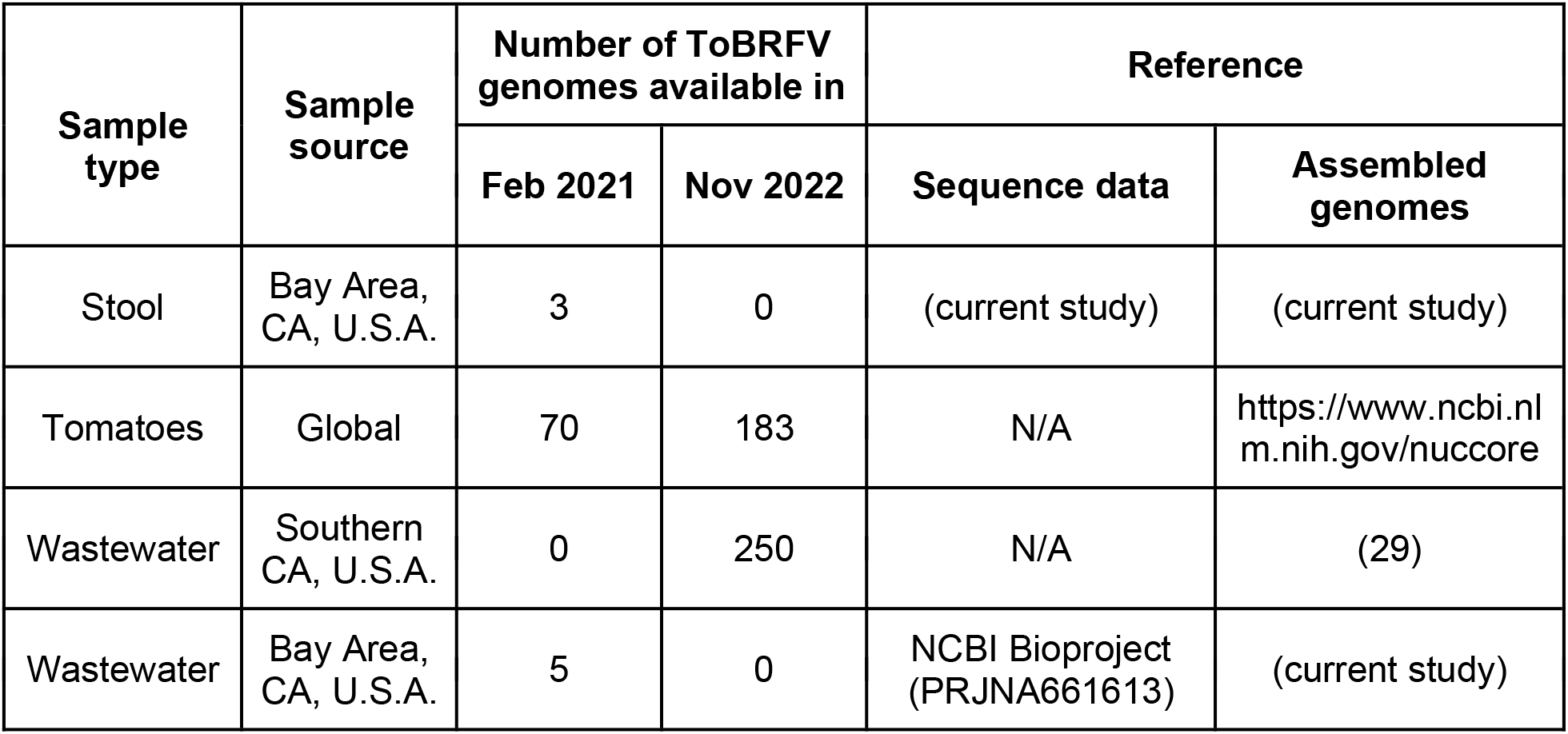
Source of genomes analyzed.

In February 2021, all near complete genomes (n = 70) of ToBRFV were downloaded from NCBI Genbank. In the same month, raw reads from the only publicly available wastewater metatranscriptomics dataset (obtained from wastewater in the Bay Area, collected between May and July 2020; Bioproject accession PRJNA661613) were also downloaded. Using these reads, five ToBRFV genomes were assembled as outlined below.

In addition to using existing sequencing data and genomes, RNA from three human stool samples obtained longitudinally from one individual were also sequenced; the first two samples were collected 10 days apart, and the third was collected 93 days after the second sample. The samples were obtained from an individual with laboratory-confirmed COVID-19 and were collected under an Institutional Review Board-approved protocol (Stanford IRB protocol #55619). Total RNA was extracted from these samples, rRNA was depleted, libraries were prepared and sequenced using NextSeq 550 as outlined in Note S1.

The following bioinformatic methods were used to assemble genomes from both the existing (from wastewater) and newly obtained (from stool) metatranscriptomics reads. Reads were trimmed with Trim Galore (version 0.4.0) using Cutadapt (version 1.8.1) (17) set to flags –q 30 and –illumina. SPAdes (version 3.14.1) set to-meta was used to assemble genomes *de novo* (17, 18). Contigs belonging to ToBRFV were classified using One Codex (19). Genes were annotated using Prodigal (version 2.6.3) set to-meta (20). If all genes were predicted on the negative strand of the contig, the entire contig was reverse complemented. The completeness of potential ToBRFV genomes was assessed using CheckV (version 1.0.1) (21) and genomes that were >90.0% complete were selected for subsequent analyses.

To assess strain diversity of ToBRFV in the longitudinal stool samples, RNA sequencing reads from stool samples were aligned to the ToBRFV reference genome (NCBI accession number NC_028478) using Bowtie (version 2.4.2) (22). The resulting bam files were used as input to inStrain (version 1.0.0) (23) to calculate popANI between genomes.

To assess abundance of ToBRFV relative to other viruses in the RNA-Seq data, reads were classified against the Viral Kraken2 database (https://benlangmead.github.io/aws-indexes/k2) (24) using default parameters. Counts from the classification were used to calculate relative abundance of viral reads.

A multiple sequence alignment of all near complete genomes of ToBRFV including genomes downloaded from NCBI Genbank in February 2021 (70 genomes) and those we assembled from wastewater and stool (8 genomes) was performed using Geneious Alignment (Geneious Prime version 2021.0.3) (25) with default settings, global alignment with free end gaps, cost similarity matrix set to 65.0%. SNPs were called from the multiple sequence alignment using SNP-Sites (version 2.5.1) (26). A phylogenetic tree was built using Geneious Tree Builder (version 2021.0.3) with default settings, Tamura-Nei genetic distance model with the neighbor-joining method. Primers and probes were designed to be specific for ToBRFV using Geneious Primer (version 3 2.3.7) (27) based on the 78 genomes we had access to in February 2021 with near default settings, requiring product size to be between 95-125 base pairs in length and primers to be based on consensus with 100.0% identity across all ToBRFV genomes. Primers and probe sequences were screened for specificity, *in silico*, using NCBI Blast.

### New genomes available in November 2022

New ToBRFV genomes became available on public databases within the last six months. An additional 113 (total 183) from NCBI (28) and 250 assembled ToBRFV genomes from a study of wastewater from Southern California (29) were downloaded (Table 1).

As Geneious alignment and tree building is computationally intensive, a phylogenetic tree of all 441 near complete genomes of ToBRFV was built using ViPTree (30), visualized and color coded by region using Iroki (31). In addition, the applicability of the primers and probes designed in this study was tested *in silico* using NCBI Blast.

### Processing of animal stool samples for RNA quantification

One stool sample was either collected from a) a single animal (cat, dog, horse, pig, rabbit) raised as a pet, b) a group of cohabiting animals of a single kind (chicken, cow, goat, mouse, sheep) from the Deer Hollow Farms (California, U.S.A.), c) a group of cohoused animals at the Deer Hollow Farms in the case of the ducks and geese, or d) from the wild (bear, deer). Samples were collected wearing gloves, using a spatula and in a sterile clinical stool collection container. Samples were transported at room temperature, aliquoted into cryovials and stored at −80°C within 12 hours from collection. Samples were further processed within a month of storage.

A single, defined solid volume of sample of each animal stool was acquired using the Integra Miltex Biopsy Punches with Plunger System (Thermo Fisher Scientific; Catalog # 12-460-410) to independent microcentrifuge tubes. 500 μl of RNALater (Ambion; Catalog # AM7023M) was added and samples were processed using a previously validated methodology (32) as follows. A stock Bovine Coronavirus (BCoV) vaccine was prepared by adding 3 mL of 1X Phosphate Buffered Saline (PBS; Fisher Scientific; Catalog # BP399-500) to one vial of lyophilized Zoetis Calf-Guard Bovine Rotavirus-Coronavirus Vaccine (Catalog # VLN 190/PCN 1931.20) to create an undiluted reagent as per manufacturer’s instructions. 10 μl of this attenuated BCoV vaccine was added to every sample as an external control and vortexed for 15 minutes. BCoV is an RNA virus that was previously found to be a reliable positive control for RNA extraction from stool (32) and helps identify instances of PCR inhibition. Samples were processed immediately after addition of the BCoV control.

### Collection and processing of human stool samples used for RNA quantification

Human stool samples were previously collected and biobanked in RNALater solution as part of Stanford Institutional Review Board-approved protocols #8903 (“Blood and Bone Marrow Grafting for Leukemia and Lymphoma”), #11062 (“Genome, Proteome and Tissue Microarray Studies in Childhood malignant and Non-Malignant Hematologic Disorders”), and #48548 (“Hematopoietic Recovery During Induction Chemotherapy in Pediatric Leukemia”). From these biobanks, 194 and 28 samples collected over the span of a year from November 2019 to October 2020 from 125 adult and 4 pediatric participants respectively were used in this study. These samples had been stored for between 1 - 12 months depending on date of collection. All samples were spiked with 10 μl of attenuated Bovine Coronavirus (BCoV) vaccine as control and processed similar to the animal stool samples.

### RNA extraction from all stool samples used for RNA quantification

RNA was extracted from these stool samples using the QIAamp Viral RNA Mini Kit (Qiagen; Catalog # 52906) as previously optimized (32). Briefly, the prepared stool samples were spun down at 10,000x *g* for 2 minutes to acquire 140 μL of clarified supernatant. RNA was extracted from this supernatant using the QIAamp Viral RNA Mini Kit (Qiagen; Catalog # 52906) as per the manufacturer’s instructions. Finally RNA was eluted in 100 μL of the elution buffer and stored in a 96-well plate at −80°C for up to 12 months. In previous work on BCoV and SARS-CoV-2 RNA (32) we found that RNA extracted using this method did not result in RT-PCR inhibitors. Therefore, we assume that samples extracted here also do not have any RT-PCR inhibitors.

### Augmenting analysis of stool with metatranscriptomic data from healthy individuals

As described below, we assessed the prevalence and abundance of MST markers in stool acquired from participants with hematologic disorders. This presented a caveat to the generalizability of our work. Therefore, we acquired metatranscriptomics data from stool samples from 10 healthy participants presented in a previous study (33). Though many human stool metatranscriptomic datasets exist, this was the most recent dataset we had access to.

### Collection and processing of wastewater samples used for RNA quantification

Settled solids were obtained from 15 wastewater treatment plants across the US (Table S2). Solids were collected from the primary clarifier, or settled from a 24 hour composited influent sample using Imhof cones. Samples were collected in sterile containers and transported to the lab. Samples from the Bay Area were processed immediately, while other samples were stored at −80°C until analysis (between 5 and 20 months).

Solids were dewatered using centrifugation and then an aliquot of the dewatered solids was set aside for dry weight analysis. Solids were then suspended in a buffer (approximately 75 mg/ml), homogenized and centrifuged. This suspension of solids in buffer was found to alleviate inhibition of RT-PCR (35). An aliquot of the supernatant was processed for total nucleic-acid extraction using Chemagic 360 (Perkin Elmer). Nucleic acid preparations from wastewater samples are known to contain PCR inhibitors that interfere with their accurate quantification using PCR-based methods. Therefore, inhibitors were removed using the OneStep PCR Inhibitor Removal Kit (Zymo research; Catalog # D6035). These methods have been published in detail (36) and step-by-step protocols are available on protocols.io (37, 38).

### Source of RNA extracted from stormwater samples used here for RNA quantification

RNA extracted from stormwater samples was derived from a previous study from our group (39). Briefly, nine stormwater samples from the Bay Area - one from Guadalupe River, Pilarcitos Creek, San Francisquito creek and San Pedro Creek, two from Stevens Creek, and three from Lobos Creek - collected between October 2018 and March 2019 were used to extract RNA (Table S3). Specifically, viruses were concentrated from 1 - 5.5 L stormwater samples using electronegative filtration using MgCl_2_. The filtration membranes were preserved in 250 μl of RNALater (Qiagen; Catalog #76104) for 5 minutes prior to storage at −80°C. Nucleic acids were extracted from the stored filtration membrane using the Qiagen DNA/RNA AllPrep PowerViral Kit using the protocol including β-mercaptoethanol and bead-beating, and stored in microcentrifuge tubes at −80°C. Previous work suggested that RT-PCR inhibitors from the samples were not co-extracted in this RNA extraction process (39). These extracts were used in the current study after 30 months of storage.

### Quantification of viral RNA sequences by ddRT-PCR

The CP gene encoding the coat protein from PMMoV, Mo gene encoding the movement protein and RdRP gene encoding the RNA-dependent RNA polymerase from ToBRFV, and M gene encoding the membrane protein from BCoV were quantified using droplet digital reverse transcription-polymerase chain reaction (ddRT-PCR). Human participants in this study were enrolled and hospitalized during the first year of the COVID-19 pandemic. We tested their stool for genes encoding the envelope (E) and a nucleocapsid (N2) protein from the SARS-CoV-2 genome as previously described (32), in order to assess occurrence of COVID-19 during hospitalization at Stanford Hospital. However, we did not find any presence of COVID-19 RNA in these samples. Sequences of the newly designed primers and probes targeting ToBRFV Mo and RdRp genes are listed in Table 2. Previously published primers and probes targeting BCoV, PMMoV and SARS-CoV-2 RNAs are listed in Table S4. We chose ddRT-PCR instead of RT-qPCR for nucleic acid detection and quantification because of its superior sensitivity and resistance to PCR inhibitors (32, 40).

**Table 2:**
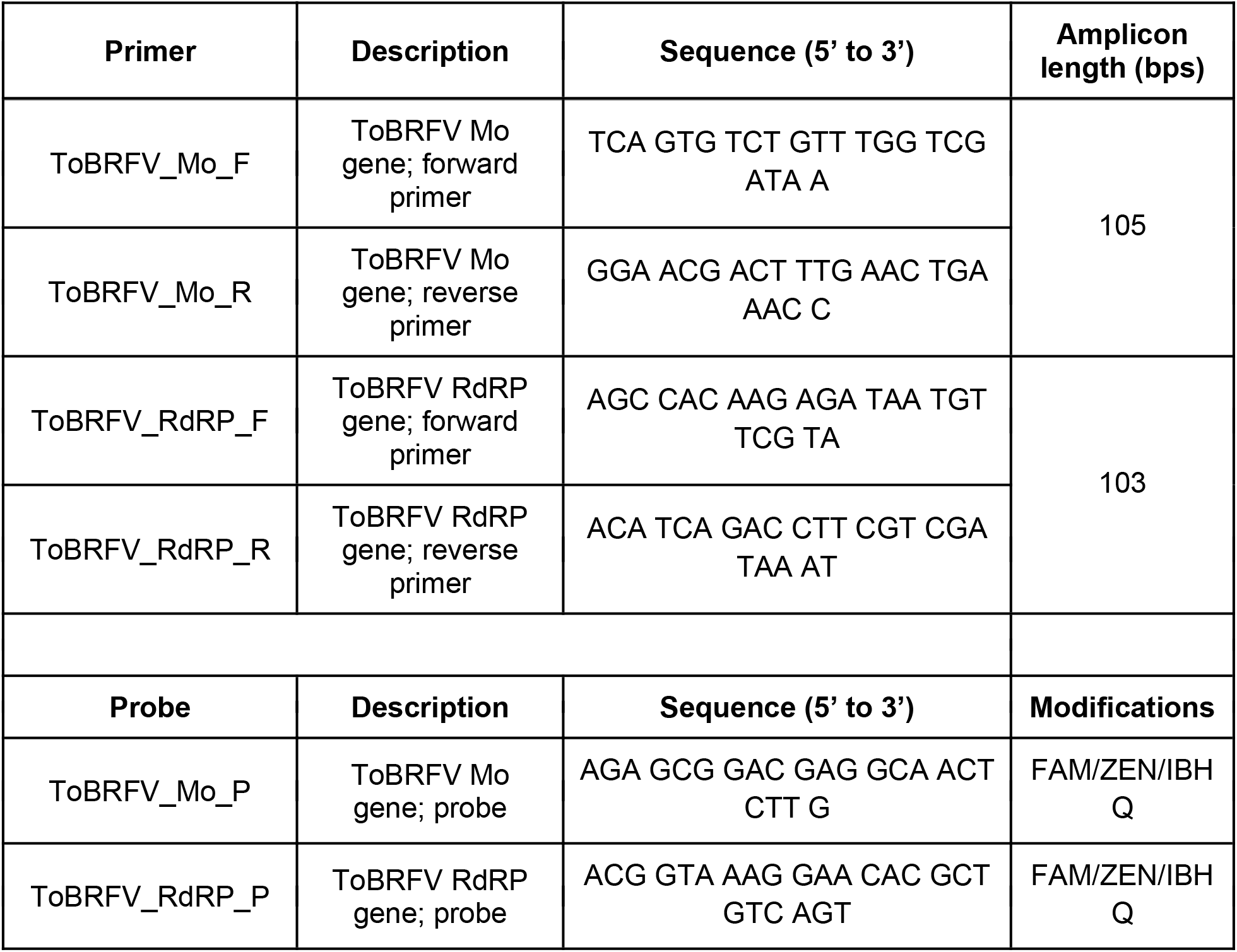
Sequences of primers and probes designed in this study to quantify ToBRFV M ands RdRP genes.

The Droplet Digital PCR Applications Guide on QX200 machines (BioRad) (41) and digital Minimum Information for Publication of Quantitative Real-Time PCR Experiments (dMIQE) guidelines (42) inform this methodology. The experimental checklist recommended by dMIQE is available at the Stanford Digital Repository (https://purl.stanford.edu/nf771cs9443). A Biomek FX liquid handler (Beckman Coulter) was used to prepare the ddRT-PCR reaction by adding 5.5 μL of eluted RNA to 5.5 μL Supermix, 2.2 μL reverse transcriptase, 1.1 μL of 300 nM Dithiothreitol (DTT), 1.1 μL of each of the 20x Custom ddPCR Assay Primer/Probe Mix (BioRad, Catalog # 10031277) and 5.5 μL of nuclease-free water (Ambion, Catalog # AM9937, Lot 2009117). The Supermix, reverse transcriptase and DTT were from the One-Step ddRT-PCR Advanced Kit for Probes (BioRad, Catalog # 1864021). A QX200 AutoDG Droplet Digital PCR System (BioRad) was used to partition the samples into droplets of roughly 1 nl using the default settings and the template was amplified using a BioRad T100 thermocycler with the following thermocycling program: 50°C for 60 min, 95°C for 10 min, 40 cycles of 94°C for 30 sec and 55°C for 1 min, followed by 1 cycle of 98°C for 10 min and 4°C for 30 min with ramp speed of 1.6°C per second at each step (43).

A multistep approach was adopted to calculate the raw RNA concentrations, as previously described (32). Every plate of ddRT-PCR assays included appropriate positive and negative controls including synthetic target genes (PMMoV CP gene PMMoV, ToBRFV Mo and RdRP genes) cloned in the pIDT vector, RNA extracted from reconstituted attenuated BCoV vaccine, water and RNA later. The signal threshold corresponding every plate was manually set between the mean positive and negative amplitudes of these controls such that the number of detected copies in the negative controls was minimal and those from the relevant positive controls most closely matched the expected RNA concentration. Next, the difference between the mean negative amplitude and the threshold amplitude in the negative control reactions was calculated and added to the mean negative amplitude for every sample on that plate. Applying this threshold yielded the raw RNA concentrations.

In order to derive the limit of blank (LoB) and limit of detection (LoD) of our assays to further process the raw RNA concentrations we adopted the following steps: a) LoB indicates the highest background RNA concentration registered from control samples that are confidently negative for the relevant gene targets. In order to determine the LoB, water, RNA later and synthetic genes discordant with the target gene (e.g. ToBRFV Mo gene is a negative control on an assay of ToBRFV RdRP gene) were assayed in duplicate. The highest RNA concentration measured in these LoB samples for each of the primer/probe sets was set as the relevant LoB. All samples that we detected an RNA concentration equal to or less than the LoB were zeroed. b) LoD is defined as the lowest concentration of RNA that can be reliably detected. To determine the LoD, duplicate serial dilution series of the synthetic target genes at the following concentrations - 1, 2, 5, 10, 100, 1000 copies/ μL of template - were assayed for the corresponding target gene (Fig. S1). The synthetic target genes were acquired from Integrated DNA Technologies and cloned in their standard backbone, pIDTSmart. These plasmids were transformed into *E. coli*, isolated using the QIAprep Spin miniprep kit (Qiagen; Catalog # 27104) and quantified using Qubit. LoD for a primer/probe set was defined as the least concentration of the standard at which both replicates had a detectable RNA concentration. All viral RNA concentrations below the LoD were zeroed.

Finally, after applying these data processing and analysis steps, the samples were assigned a final viral RNA concentration in copies/ μL of template. Eluate refers to the 100 μL of sample acquired from the RNA extraction. Viral RNA concentrations from animal and human stool samples are expressed as copies/ μL of template, from wastewater samples as copies/ g of wastewater, and from stormwater samples as copies/ liter of stormwater.

In the case of all non-human animal stool, wastewater and stormwater samples, RNA was quantified using singleplex reactions. For the human stool samples, which were limited in quantity, the detection of the BCoV M gene and PMMoV CP gene were multiplexed with the detection of the SARS-CoV-2 E and N2 genes using orthogonal fluorescent probes. After extensive optimization (outlined in the Note S1), we paired the detection of the E gene (SARS-CoV-2) with the CP gene (PMMoV), and N2 gene (SARS-CoV-2) with the M gene (BCoV) in two independent reactions using the carboxyfluorescein (FAM) and hexachlorofluorescein (HEX) fluors, respectively.

### Data analysis and generation of plots

Data was analyzed using RStudio (ver 1.2.5042), using packages cowplot (ver 1.1.1), dplyr (ver 1.0.8), eulerr (ver 6.1.1), ggplot2 (ver 3.3.6), and UpSetR (ver 1.4.0).

### Data availability

Newly generated genomes and raw sequencing reads from stool samples are available on NCBI’s Sequence Read Archive (SRA) database at accession #PRJNA917455. All other relevant data are included in this manuscript and available through the Stanford Digital Repository (https://purl.stanford.edu/nf771cs9443).

## Results and discussion

### ToBRFV is widely prevalent and abundant in sequence data from stool and wastewater samples

Tracking the presence of human feces in the environment, and the identification of internal controls for the processing of stool and wastewater samples requires marker genes that are: a) prevalent - consistently present across samples, and b) abundant - available in high enough concentration for reliable detection. crAssphage (12) presents one such DNA-based marker and PMMoV is another RNA-based marker (13). We sought to test whether RNA-based markers from ToBRFV also meet these criteria.

We isolated and sequenced RNA from three longitudinal stool samples from one human participant, who had tested positive for SARS-CoV-2. In parallel, we acquired publicly available transcriptomics data from five wastewater samples that had been collected and sequenced from the Bay Area (44). Using these sequence data from eight samples, we identified all represented RNA viruses and their relative abundances (Fig. 1). The tomato brown rugose fruit virus (ToBRFV) was the most widely prevalent RNA virus, present in all five wastewater samples and three stool samples. It was detected at very low relative abundance (0.077% of viral reads) in one of the stool samples during the time of active SARS-CoV-2 infection, in which 99.9% of reads belonged to SARS-CoV-2. In the other seven samples where it is detected, it is the only viral RNA that is consistently over 10.0% in relative abundance of viral reads, often making up over 50.0% of the reads. Notably, the relative abundance of ToBRFV was consistently greater than PMMoV, which is a well established microbial source tracking marker and known to be highly abundant in wastewater (8). This is consistent with reports from studies carried out prior to (44), and in parallel with (29, 45) ours, that also show that ToBRFV is a highly prevalent virus in wastewater.

**Fig. 1:**
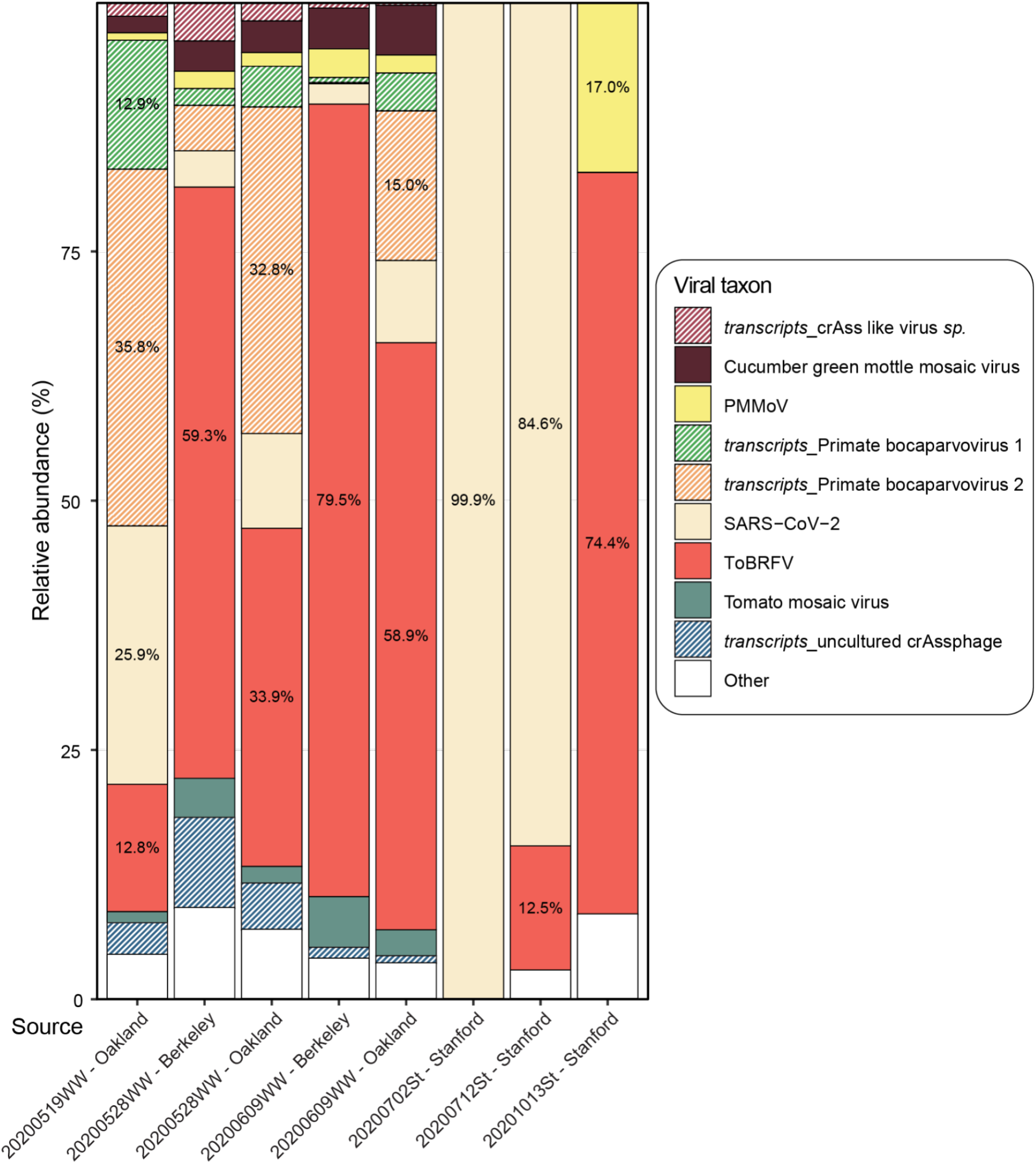
Relative abundance of viral RNA from sequencing wastewater and stool samples. The x-axis represents the source of the eight sequencing datasets analyzed here. Five wastewater samples are marked by the date of collection in ‘YYYYMMDD’ format followed by “WW” and the location of collection. The three stool samples are marked by the date of collection in ‘YYYYMMDD’ format followed by “St” and the location of collection. The y-axis indicates the relative abundance of each taxon. The color scheme represents specific taxon as light brown for crAss like virus *sp*., dark brown for cucumber green mottle mosaic virus, yellow for PMMoV, green for primate bocaparvovirus 1, orange for primate bocaparvovirus 2, cream for SARS-CoV-2, salmon for ToBRFV, light blue for tomato mosaic virus, dark blue for uncultured crAssphage, and white for other minor taxa. Patterned bars highlight sequence reads from transcripts from taxa that have DNA genomes. Taxa with >10.0% relative abundance also list the percentage of abundance in the histogram.

### Novel ToBRFV genomes and sequence analysis reveal suitable RNA-borne marker genes

Having identified that ToBRFV is a prevalent and abundant RNA virus in sequence data, we next set out to identify genomic regions suitable as a target for primer/probe for its reliable molecular detection.

In February 2021, at the start of this study, only 70 near complete ToBRFV genomes were known. 50 of these were from the Netherlands. None had been sequenced from human stool or wastewater samples, and only one sequence was derived from the U.S.A. In order to ensure that the assay we developed was universal, we first decided to augment the number of ToBRFV genomes and the diversity of its sources. Therefore, we assembled near complete genomes of ToBRFV using sequence data generated in this study from stool samples and using existing data from wastewater samples (44), both collected in the Bay Area. The eight newly assembled genomes had a mean completeness of 98.8% (range 93.6% - 100.0%; median 99.4%) (Table S5). The longitudinally acquired stool samples yielded ToBRFV genomes with SNPs in 27 positions, suggesting possible strain variation over time. Looking more broadly, across all 78 near complete ToBRFV genomes, we identified 2,808 positions containing SNPs (across an average contig length of 6,366 bp), and the 12 North American strains form their own, distinct cluster (Fig. 2B).

**Fig. 2:**
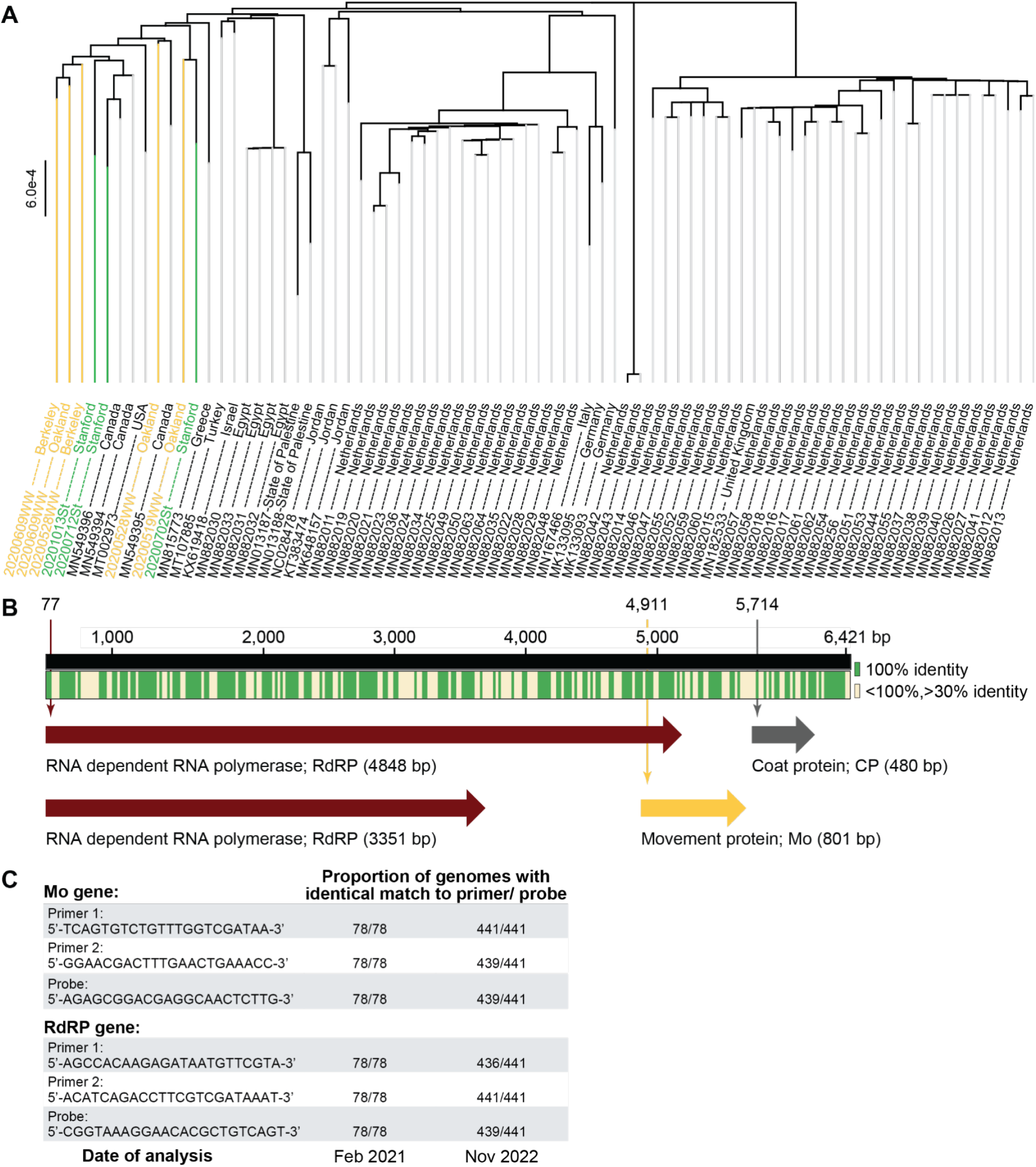
Analysis of newly assembled ToBRFV genomes and generation of primer/probe sets for ddRT-PCR. (A) Phylogenetic tree of 78 near complete genomes of ToBRFV, including eight genomes generated in the current study from wastewater and stool. All genomes are listed by their NCBI accession number and source location. 70 pre-existing genomes are listed in black font, five genomes derived from wastewater samples are listed in yellow, and three from stool samples in green. (B) Summary of multiple sequence alignment and gene annotation across the 78 ToBRFV genomes. Green indicates regions that are 100.0% conserved across all genomes, while cream marks those that are greater than 30.0% but less than 100.0% conserved. Two variants of the RNA dependent RNA polymerase (RdRP) encoding gene are found at 77 bp and are of either 4,848 bp or 3,351 bp in size. The movement protein (Mo) encoding gene is found at 4,911 bp and is 801 bp in size. The coat protein (CP) encoding gene is found at 5,714 bp and is 480 bp in size. Genomic locations are based on genome ID NC_028478. Sequences of primer/probe sets generated in the current study, in Feb 2021, aimed at targeting the Mo and RdRP genes across all known genomes. Since the number of known genomes grew from Feb 2021 to Nov 2022, the final column indicates the proportion of the 441 current genomes bearing sequences identical to the designed primer/probe sets.

Multiple sequence analysis across all 78 ToBRFV genomes highlights regions that are 100.0% conserved (Fig. 2C). Among these, gene annotation reveals a) two variants of the RNA dependent RNA polymerase (RdRP) encoding gene at 2,700 bp on the chromosome, that differ by whether an internal stop codon is read through (size 3,351 bp or 4,848 bp), b) the movement protein (Mo) encoding gene (size 480 bp) at 5,166 bp, and c) the coat protein (CP) encoding gene (size 801 bp) at 5,166 bp (Fig. 2C). Among these we designed primer/probe sets targeting the 5’ end of the RdRP gene, and the Mo gene. We were unable to identify a suitable primer set for the CP gene for ddRT-PCR. Notably, the primer/probe sets designed here (Table 2) were conserved across all 78 genomes (Fig. 2D).

Within the last six months, the number of near complete ToBRFV genomes have increased to 441 (Table 1), with additional genomes from Belgium, France, Mexico, Switzerland, and the U.S.A. (summarized in Fig. 2b). Therefore, we repeated the phylogenetic analysis of the novel genomes generated in the current study in the context of all 441 currently known genomes (Fig. S3). Again, we find that the genomes derived from North American cluster distinctly. Finally, we analyzed whether the primer/probe sets proposed here continue to be universal and found that the oligonucleotides targeting Mo are a perfect sequence match in 439/441 genomes, while those targeting RdRP are a perfect match in 436/441 genomes (Fig. 2D).

### ToBRFV targeting primer/probe sets have low LoB and LoD

Having newly designed primer/probe sets targeting the Mo and RdRP genes in ToBRFV, we aimed to validate these oligonucleotides and establish the limits of their reliable utility.

To this end, we acquired synthetic DNA constructs featuring regions of the ToBRFV Mo and RdRP genes targeted by hydrolysis-probe RT-PCR assays from Integrated RNA Technologies (IDT) cloned into the pIDT plasmid. We also acquired a similar plasmid containing the PMMoV CP gene. Using ddRT-PCR, we assayed a dilution series of these synthetic plasmid constructs at 1, 2, 5, 10, 100, 1000 copies / μL of template in triplicate and found that all the primer/probe sets detected the target gene at all concentrations (Fig. S4). Next, we focused our attention on the negative controls included in the assays to identify the limit of detection for each primer/probe set. The negative controls included two no template controls, water and RNALater, and two mismatched controls that were the synthetic pIDT plasmids bearing targets orthogonal to the primer/probe sets. Therefore, theoretically, all the negative controls would have no detectable gene target. For each primer/probe set, among the negative controls, we identified the highest concentration of target detected and set this value as the limit of blank (LoB). This means any concentration below −0.552 log_10_ copies/ μL of template for primer/probe set targeting PMMoV CP gene, −0.590 log_10_ copies/ μL of template for ToBRFV Mo gene, and 0.407 log_10_ copies/ μL of template for ToBRFV RdRP gene is not reliable (Fig. S4). After converting all concentrations of gene targets below the LoB to zero, we focused our attention on the triplicate dilution series to identify the lowest concentration of template at which all three reactions had a detectable target concentration (Fig. S4). We set this concentration as the limit of detection (LoD), i.e. the least concentration at which a gene target can be reliably detected. The LoD for the primer/probe set targeting PMMoV CP gene was 1 copies/ μL of template, for ToBRFV Mo gene was 5 copies/ μL of template, and for ToBRFV RdRP gene was 5 copies/ μL of template. All gene target concentrations below the LoD were set to zero.

### ToBRFV is not detected in stool from non-human animals

MST targets should be specific, meaning they are mostly absent in stool from other common animals. Therefore, having established that our primer/probe sets are functional, we tested them against stool collected from 14 different animals including wild bear and deer, chicken, cow, duck, goose, goat and sheep from a farm, horse and pig from a barn, a household cat, dog and rabbit, and laboratory mice. Notably, these animals are rather diverse and are fed a wide variety of foods. While RNA extracted from all of these animal samples had a detectable concentration of the M gene target from the spiked-in BCoV used as a control, none of them had RNA containing either the PMMoV CP gene or the ToBRFV RdRP gene. The ToBRFV Mo gene was detectable only in the sample derived from the domesticated cat, perhaps due to inclusion of tomatoes in its processed kibble or cross contamination of diet with its human cohabitant. Therefore, all three sets of primer/probe to detect RNA from PMMoV and ToBRFV are not detected in most animal feces, except for the ToBRFV Mo gene in a cat, indicating they are specific for human stool.

### Description of participants who provided human stool samples used for RNA quantification

Analyzing sequence information from three stool samples collected from one human participant revealed ToBRFV to be abundantly present. To further test the sensitivity of the assays to human stool, we relied on a stool biobank including 194 stool samples from 125 adults and 28 samples from four children, all of whom were undergoing hematopoietic cell transplantation (HCT), cell therapy (CAR-T) or induction chemotherapy for the treatment of underlying hematologic disorders.

Of the adult participants, 79 are male, 45 are female and 1 participant did not provide information on their sex. The median age of the adult participants is 60 years (range 19 - 82 years) and pediatric participants is 6 years (range 3 - 16 years). Among the adult participants, 61.6% self-identify as White. Age, race and ethnicity information on pediatric participants are withheld since these can be used to identify the participants. Timeline of stool collection is summarized in Fig. S1. Demographic information is summarized in Fig. S2 and Table S1.

### ToBRFV is more prevalent in human stool samples compared to PMMoV

220 out of 222 RNA extracts derived from 129 participants had detectable BCoV RNA. This suggests that two of the RNA extractions failed; those samples are therefore excluded from further analysis, altering our study cohort size to 127 (123 adult; 4 pediatric). Among the remaining samples, 126/220 (57.3%) of stool samples had detectable levels of the PMMoV CP gene, while 143/220 (65.0%) had the ToBRFV Mo gene and 108/220 (49.1%) had the ToBRFV RdRP gene; ToBRFV Mo gene was the most prevalent target gene. This prevalence varied in the two patient cohorts (Fig. 4A); 127/192 (66.2%) stool samples from adult participants had detectable amounts of the ToBRFV Mo gene, more than in the case of PMMoV CP gene (103/192; 54.7%), but only 16/28 (57.1%) stool samples from pediatric patients had detectable amounts of the ToBRFV Mo gene, fewer than in the case of PMMoV CP gene (23/28; 82.1%).

**Fig. 3:**
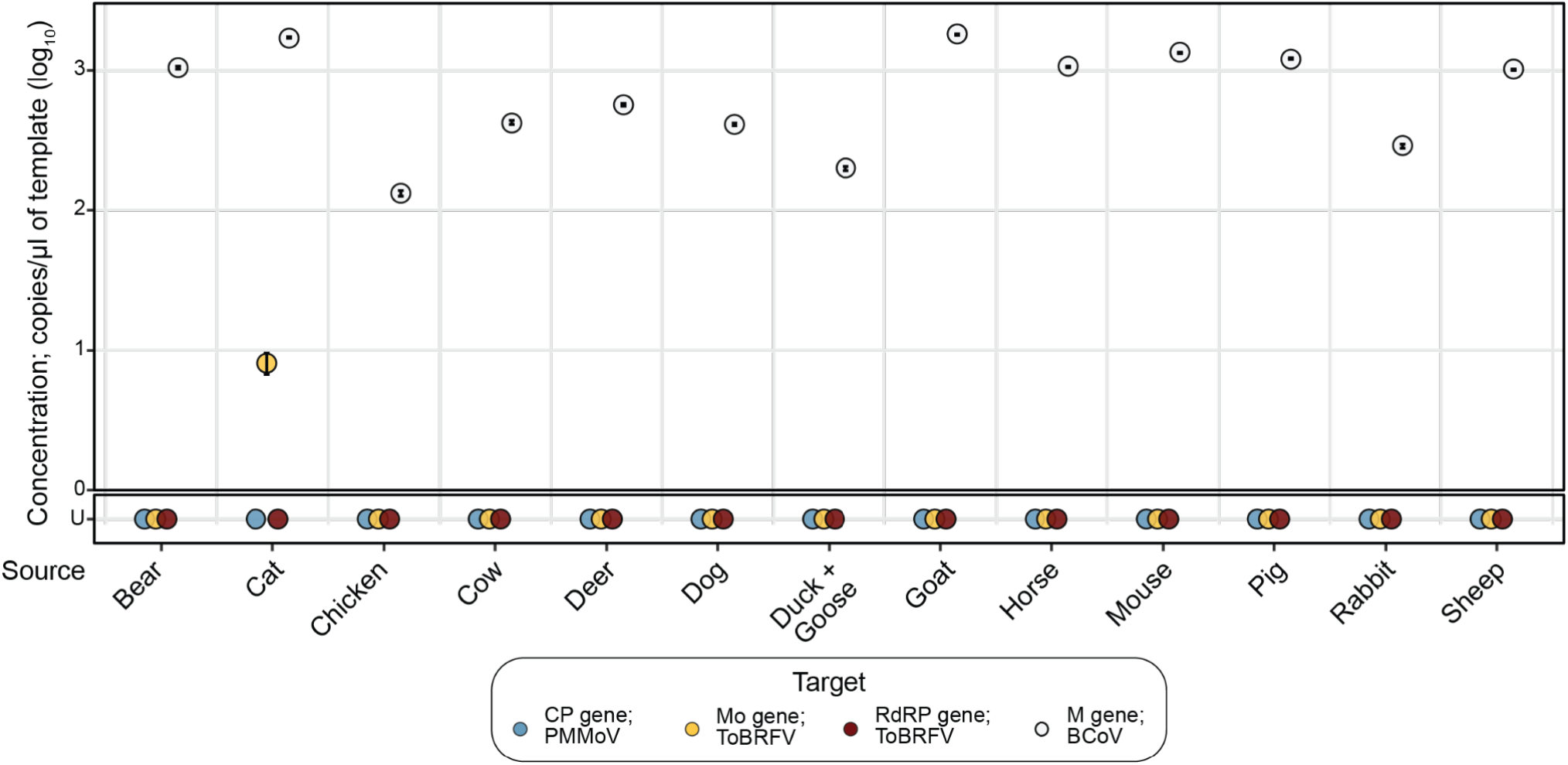
Concentrations of PMMoV and ToBRFV target genes in animal stool samples. Dot plot marking the concentrations of the PMMoV CP (blue), ToBRFV Mo (yellow) and RdRP (red) genes. Concentrations of BCoV M gene, used as a control, are marked by white dots. Error bars marking the standard deviation are plotted along with the dots, and are mostly subsumed within the dot. The x-axis lists 13 samples from 14 different animals, where a single sample is derived from cohoused ducks and geese. The y-axis lists concentrations of the genes in log_10_ copies/ μl of template; U stands for “Undetermined” and marks samples with no detectable gene

**Fig.4:**
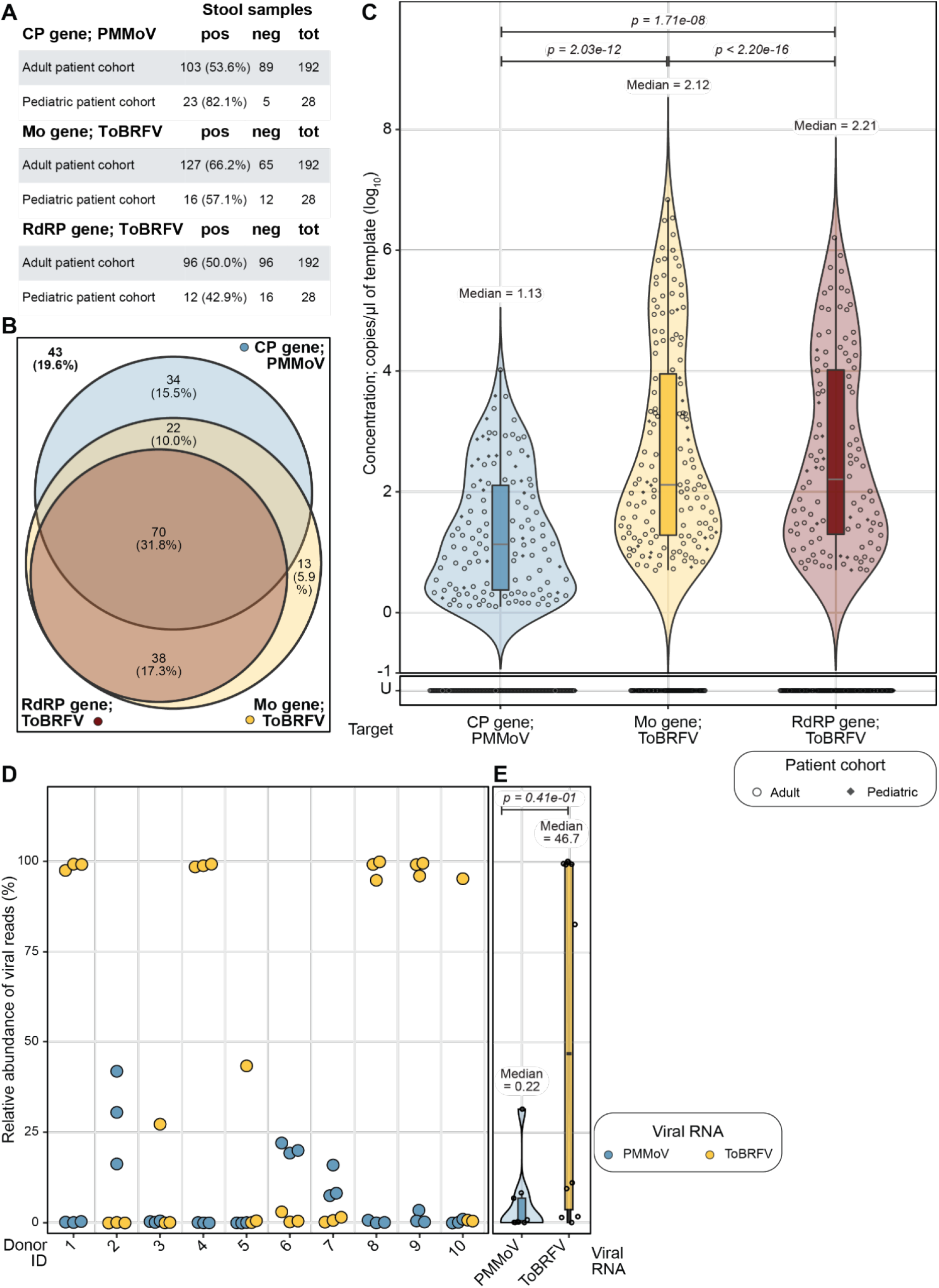
Prevalence of PMMoV and ToBRFV target genes in human stool samples. (A) Tabular summary of detection of the three gene targets in samples from adult and pediatric cohorts. The first column lists the name of the target gene and cohort, followed by the number and percent of samples that were positive (pos) or number of samples that were negative (neg) for that target gene, and the total number of samples tested (tot). (B) Venn diagram summarizing the detection of the PMMoV CP gene (blue), and ToBRFV Mo (yellow) and RdRP (red) genes across 220 human stool samples. In 34 (15.5%) we only detected the PMMoV CP gene, while 13 (5.9%) only the ToBRFV Mo gene. In 38 (17.3%) samples we detected both ToBRFV target genes, while 22 (10.0%) had both the PMMoV CP gene and ToBRFV Mo gene. In 70 (31.8%) of the samples we detected all three gene targets, while 43 (19.6%) had none of them. (C) Dot plot marking the concentrations of PMMoV CP (blue), ToBRFV Mo (red) and RdRP (yellow) genes, with violin and box plots summarizing their distributions, in RNA extracted from stool samples collected from humans. The x-axis marks the target genes, and the y-axis lists their concentrations in log_10_ copies/ μl of template; U stands for “Undetermined” and marks samples with no detectable gene target above LoB. Adult samples are marked by unfilled circles and pediatric samples are marked by a filled diamond. The concentration of PMMoV CP gene is a median of 1.13 with a standard deviation of 1.00 and IQR of 1.74 log_10_ copies/ μl, ToBRFV Mo gene is a median of 2.12 with a standard deviation of 1.69 and IQR of 2.67 log_10_ copies/ μl, and ToBRFV RdRP gene is a median of 2.20 with a standard deviation of 1.56 and IQR of 2.72 log_10_ copies/ μl. *p* values derived from paired Wilcoxon signed-rank tests with continuity correction and excluding samples with undetermined concentration across all combinations of the three gene targets are listed at the top of the plot. (D) Dot plot marking the relative abundance of viral reads of PMMoV (blue) and ToBRFV (yellow) from previously published metatranscriptomics data derived from healthy stool samples. The x-axis lists the 10 donors who provided samples, and each sample provided RNA sequences in biological triplets; each dot denotes a single replicate. The y-axis lists relative abundance in percent. (E) Dot plot summarizing data from panel D, now including violin and box plots to highlight distribution of viral RNA concentrations and associated statistics. The x-axis marks the target viral RNA, and the y-axis lists their relative abundance in percent. Dots represent the average of data from three biological replicates. PMMoV (blue) is present at a median relative abundance of 0.217% with a standard deviation of 9.83% and IQR of 5.19%, ToBRFV (yellow) is present at a median relative abundance of 46.7 with a standard deviation of 48.5% and IQR of 95.4%. *p* value derived from a Wilcoxon signed-rank test of pairwise differences in relative abundance with continuity correction and excluding samples with undetermined concentration is listed at the top of the plot.

In analyzing the prevalence of the three gene targets of interest in the stool samples, we detected all three gene targets in 70 (31.8%) of the samples, while we detected none of the three gene targets in 43 (19.6%) (Fig. 4B, Fig. S5). Notably, in 34 (15.5%) of the samples, we only detected the PMMoV CP gene, and in 13 (5.9%), we only detected the ToBRFV Mo gene. In all samples that we detected the ToBRFV RdRP gene we also detected the ToBRFV Mo gene. This analysis suggests that while the ToBRFV Mo gene is the most prevalent RNA-based marker of human stool, combining this with the detection of the PMMoV CP gene will provide the most coverage of more than 80.0% of stool samples.

Next we analyzed the abundance of each of these gene targets in stool samples. The median detected concentration of the PMMoV CP gene is lower than the ToBRFV Mo gene (1.13 log_10_ copies/ μl vs. 2.12 log_10_ copies/ μl, Wilcoxon signed rank test *p* = 2.03e-12) and the ToBRFV RdRP gene (1.13 log_10_ copies/ μl vs. 2.21 log_10_ copies/ μl, *p* = 1.17e-8; Fig. 4C). These stool samples were derived from participants undergoing different treatments for underlying hematologic disorders. Therefore, we investigated whether the nature of treatment was a confounding factor. Here, again, we find that the median abundances of both target genes from ToBRFV are higher than the PMMoV CP gene, even when the samples are separated by treatment cohort (Fig. S6A). Further, a paired comparison of target gene abundances validates the previous observation that all samples that tested positive for the ToBRFV RdRP gene also tested positive for the ToBRFV Mo gene (Fig. S6B).

While the concentration of the various gene targets has so far been reported in copies per μl of template, we recognize that studies also measure molecular targets in units per g dry weight of stool sample. Therefore, we chose five samples per cohort at random, dried two biopsy punches from each sample and found the mean percent dry weight of the samples from adults undergoing HCT treatment as 23.6% (range 18.2 - 33.9%), CAR-T treatment as 27.5% (range 22.2 - 31.6%), and those of pediatric patients undergoing induction chemotherapy as 32.4% (range 23.8 - 40.3%). We used the average percent dry weight to convert gene target concentrations to copies per gram (g) dry weight of stool samples in Fig. S6C. In brief, the median concentrations of ToBRFV RdRP and Mo genes were 6.45 and 6.32 log_10_ copies/ g dry weight of stool, the PMMoV CP gene was 5.36 log_10_ copies/ g (Fig. S6C).

To determine if our findings are generalizable to applications beyond a cohort of patients, we looked at an alternate dataset recently generated in our group that sequenced RNA from stool collected and frozen from 10 healthy individuals in triplicate (33). In this dataset also, the relative abundance of ToBRFV was consistently greater than PMMoV, as reflected by their median relative abundance of 46.7 vs. 0.22% viral RNA reads (Fig. 4D). Taken together, the abundance of ToBRFV is greater than PMMoV in human stool samples, and ToBRFV Mo gene may thus be a preferable MST marker compared to the the PMMoV CP gene.

### ToBRFV Mo gene is prevalent and abundant in wastewater samples

Wastewater is a complex matrix containing human stool and other biological excretions, in addition to food waste, industrial wastes, and infiltrated stormwater in some cases. We next validate the molecular detection test developed here for testing this sample type. We acquired wastewater solids samples from 15 cities in the U.S.A, extracted RNA and assayed these for the presence and abundance of the gene targets of interest.

The extracted RNA from Wisconsin failed to have detectable amounts of any of the gene targets; this matches unpublished data generated using this sample by a different group and is excluded from further analysis reducing our sample size to 14. 13 of these samples have more ToBRFV Mo gene than the other two molecular markers, with the sample from New York being the exception, having PMMoV CP gene in the highest concentration (Fig. 5A). Looking at the data in aggregate, the samples have a median concentration of 10.5 log_10_ copies/ g dry weight of wastewater solids with a standard deviation of 0.67 and IQR of 0.26 log_10_ copies/ g of the ToBRFV Mo gene, followed by 9.81 log_10_ copies/ g, standard deviation of 0.60 and IQR of 0.36 log_10_ copies/ g of the ToBRFV RdRP gene, and 9.49 log_10_ copies/ g, standard deviation of 0.46 and IQR of 0.74 log_10_ copies/ g of the PMMoV CP gene. Pair-wise comparison of gene target concentrations across samples using the Wilcoxon signed-rank test reveals that the increased detection of the ToBRFV Mo gene is statistically significant in comparison to the PMMoV CP gene (*p* = 1.37e-3) and the ToBRFV RdRP gene (*p* = 1.10e-3).

**Fig. 5:**
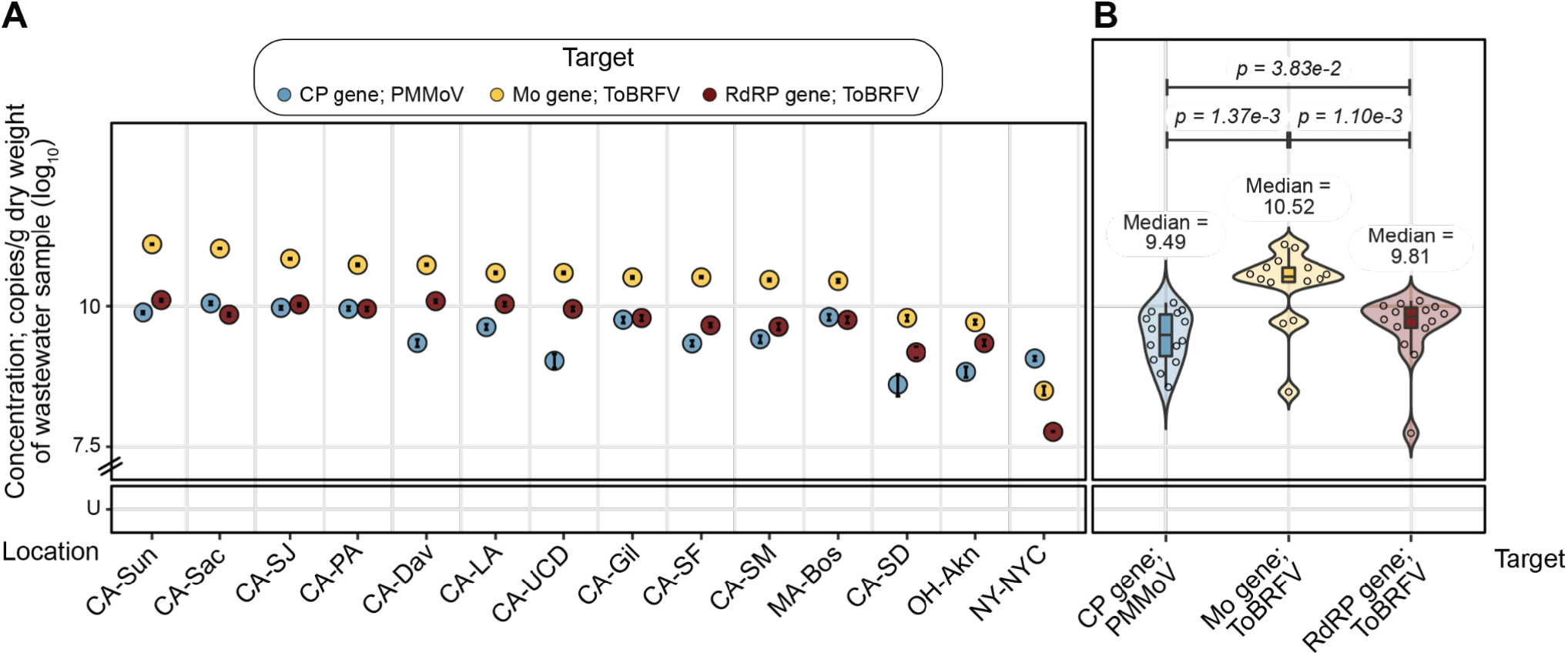
Concentrations of PMMoV and ToBRFV target genes in wastewater samples from across the U.S.A. (A) Dot plot marking the concentrations of PMMoV CP (blue), ToBRFV Mo (yellow) and ToBRFV RDRP (red) genes across samples. Error bars marking the standard deviation are plotted along with the dots, and are mostly subsumed within the dot. The x-axis lists the 15 cities from where samples were sourced in decreasing concentration of the Mo gene; abbreviations for state and cities are expanded in Table S2. The y-axis lists concentrations of the genes in log_10_ copies/ g dry weight. (B) Dot plot marking the concentrations of PMMoV CP (blue), ToBRFV Mo (red) and RdRP (yellow) genes, with violin and box plots summarizing their distributions, in RNA extracted from wastewater samples collected from across the U.S.A. The <-axis marks the target genes, and the y-axis lists their concentrations in log_10_ copies/ g of vastewater; U stands for “Undetermined” and marks samples with no detectable gene target above LoB. The PMMoV CP gene has a median of 9.49 with a standard deviation of 0.46 and QR of 0.74 log 10 copies/ g dry weight of wastewater sample, ToBRFV Mo gene has a median of 10.5 with a standard deviation of 0.67 and IQR of 0.26 log_10_ copies/ g, and ToBRFV RdRP gene has a median of 9.81 with a standard deviation of 0.60 and IQR of 0.36 log_10_ copies/ g. *p* values derived from paired Wilcoxon signed-rank tests with continuity correction across all combinations of the three gene targets are listed at the top of the plot. U stands for “Undetermined” and marks samples with no detectable gene target above LoB.

### ToBRFV Mo gene matches crAssphage ORF000024 as an indicator of fecal contamination of stormwater

crAssphage ORF000024 is a well established human-associated microbial source tracking marker (12). We compared concentrations of PMMoV and ToBRFV RNA targets to those of this crAssphage DNA target in stormwater draining from urbanized watersheds in the Bay Area. crAssphage ORF000024 was previously quantified in these samples and is reported in Graham *et. al*. (39).

We found that in the nine stormwater samples, crAssphage ORF000024 had the highest median concentration of 4.65 with a standard deviation of 0.56 and IQR of 0.66 log_10_ copies/ liter of stormwater, followed by the ToBRFV RdRP gene with a median of 3.48, standard deviation of 0.97 and IQR of 1.24 log_10_ copies/ liter of stormwater, ToBRFV Mo gene with a median of 3.34, standard deviation of 0.99 and IQR of 1.36 log_10_ copies/ liter of stormwater, and finally the PMMoV CP gene with a median of 3.02, standard deviation of 0.54 and IQR of 0.44 log_10_ copies/ liter of stormwater (Fig. S7). Pair-wise comparison of gene target concentrations across samples using the Wilcoxon signed-rank test reveals that differences in concentrations are not statistically significant and gene targets are similarly abundant. The concentration of gene targets in each of the samples is presented in Fig. S7. Notably, the ToBRFV Mo gene is detected in as many samples (6/9) as crAssphage ORF000024 (Fig. 6). This result suggests using an RNA based marker from ToBRFV to detect human stool contamination of storm water may be as useful as using the DNA marker from crAssphage ORF000024.

**Fig. 6:**
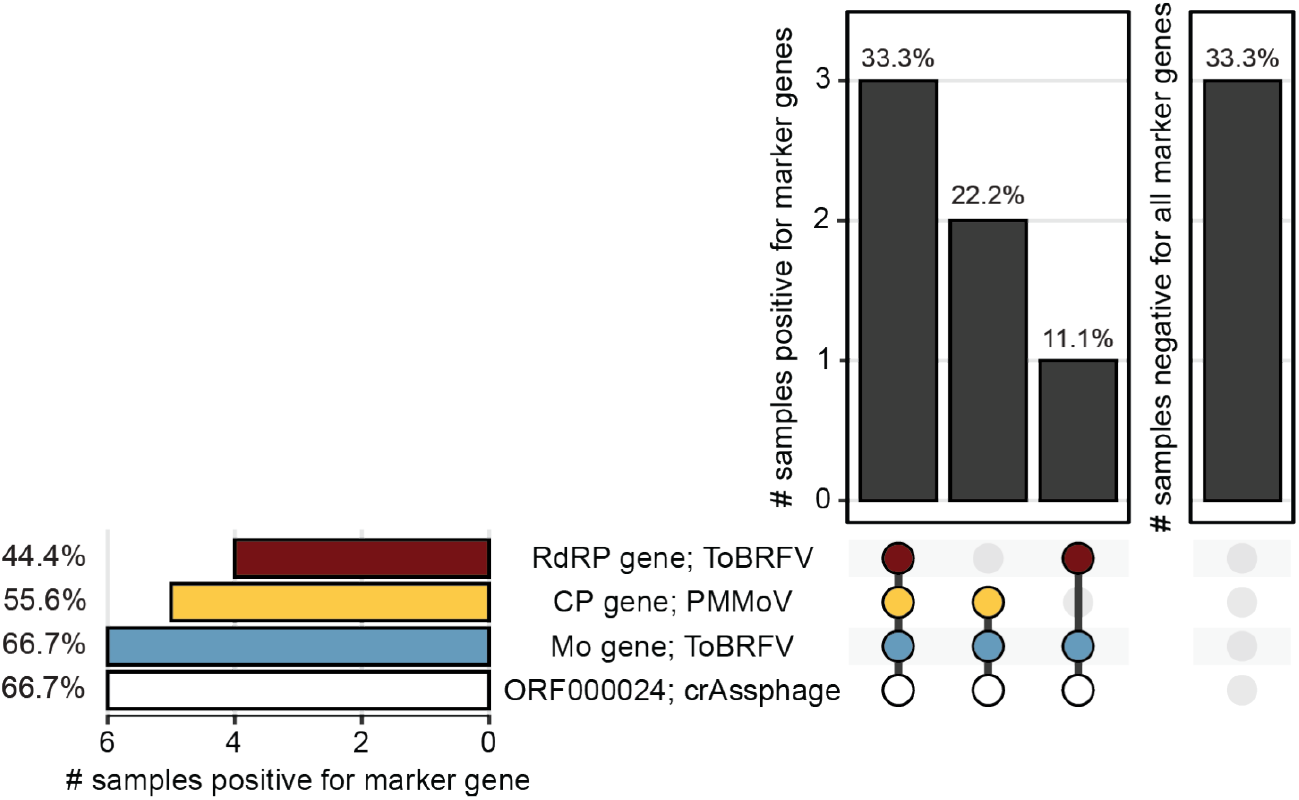
Prevalence of PMMoV, ToBRFV and crAssphage target genes in stormwater samples from across California. UpSet plot summarizing the number of stormwater samples (total n = 9) that are either positive for multiple marker genes (left), or negative for all marker genes (right) in the vertical bar plots. Marker genes are listed under the plots, with colored dots representing positive presence and grey dots representing absence. Marker genes present in samples represented in the vertical bar are also connected by a thick line. Prevalence of independent marker genes are also summarized in the horizontal bar plot. All bars list data in percentage units. PMMoV CP gene is marked by blue, ToBRFV Mo and RdRP genes by yellow and red respectively, and crAssphage ORF000024 by white. Data on crAssphage are derived from a previous study (39).

## Conclusions and limitations

In this study, we generate eight nearly complete genomes of ToBRFV from wastewater and stool from the Bay Area. We catalog SNPs in all existing genomes, including in those that we assembled here, and note variations in viral genomes isolated from the same individual over ~ 100 days. We then went on to identify two sets of primers and probes that can universally detect ToBRFV across the world.

Assays developed using these primer, probe sequences are sensitive and specific for human stool and wastewater as it was present in a wide range of wastewaters and stool samples, and not present in any tested animal stool aside from one cat. Like the established viral MST target PMMoV (8), the ToBRFV target is derived from the genome of a plant virus likely present in the human gut owing to dietary intake of diseased plants. Concentrations of ToBRFV Mo and RdRP gene targets were as high or higher than PMMoV CP gene in wastewaters and stormwater known to contain sewage. The ToBRFV targets’ high concentrations in wastewater, as well as in human stool samples, suggests that they may be useful as endogenous fecal strength controls for wastewater-based epidemiology applications (46), as well as an endogenous positive extraction control during nucleic acid extractions in studies seeking to quantify rare infectious disease targets (8).

There are several limitations to this work. First, the specificity of the ToBRFV Mo and RdRP gene targets was tested using just one representative sample of various non-human, animal stools. Additional work to test more animal stool samples would be helpful to further characterize the assays’ specificity for human stool. Second, the sensitivities of the various assays were tested using only human stool samples from individuals residing in the Bay Area. It is possible that the distribution of the targets in individuals from other locations may differ from those studied here and more work is encouraged to document the ToBRFV prevalence and abundance in samples globally. Third, we assayed wastewater solids sampled from around the U.S.A, from New York to California, and they contained high concentrations of the ToBRFV targets. Further work from samples around the world will be valuable to testing the generalizability of the assays. Notably, the presence of ToBRFV genomes from this study and others collected from many countries reassures us that ToBRFV is likely to be a universal global MST marker. Finally, As more ToBRFV genomes become available, it will be important to test whether the primers and probes developed herein continue to overlap with conserved regions of the genomes.

## Acknowledgements

This work was supported by a ChemH-IMA grant (to A.S.B.), a gift from the CDC-Foundation (to A.B.B.), Stanford MCHRI and NIH T32 DK098132 (C.J.S.), DGE-1656518 and T32GM007276 (to D.T.S.), and the Stanford Dean’s Postdoctoral Fellowship (to A.N.). A.S.B. laboratory is supported by NIH R01 AI148623 and R01 AI143757. We acknowledge Aaron Behr, Alvin Han, David Miklos, David Solow-Cordero, Dhananjay Wagh, Jennifer Estes, Isabel Delwel, Luisa Jiminez, Said Attiya, Sopheak Sim, and Summer Vance for technical assistance, and sharing of samples and data for use in this study. A.N., B.J.F, M.W. and A.B.B., are co-inventors on a U.S. provisional patent application #63/387,657 that has been filed and relates to the methods presented in this manuscript. The other authors declare no competing interests. This study was performed on the ancestral and unceded lands of the Muwekma Ohlone people. We pay our respects to them and their Elders, past and present, and are grateful for the opportunity to live and work here.

